# Genetic response to light and carbon source variations in *Trichoderma harzianum*: The key regulatory roles of *env1*, *cre1*, and *blr2*

**DOI:** 10.1101/2025.05.30.657067

**Authors:** Rafaela Rossi Rosolen, Maria Augusta Crivelente Horta, Danilo Augusto Sforca, Anete Pereira de Souza

## Abstract

In filamentous fungi, light plays a key role in regulating physiological processes such as growth, conidiation, secondary metabolism, and the expression of hydrolytic enzymes. The processes that depend on light are controlled by photoreceptors, including BLR1, BLR2, and ENV1, as well as by signaling pathways involving heterotrimeric G-proteins and cyclic adenosine monophosphate (cAMP). *Trichoderma harzianum* is a promising candidate for biotechnological use and is able to promote hydrolytic reactions under biomass degradation conditions. However, the genetic mechanisms underlying its response to light remain poorly understood, especially under degradative conditions. This study aimed to assess the expression of carbohydrate-active enzymes (CAZymes), transcription factors (TFs), and signaling pathway proteins under different light conditions and carbon sources. The results revealed distinct patterns of relative gene expression influenced by these environmental factors, highlighting the complex regulatory mechanisms at play in *T. harzianum*. Moreover, our results suggested that *env1*, *cre1*, and *blr2* are critical for adjusting to different light conditions and carbon sources. This highlights the importance of both factors in regulating gene expression and supporting metabolic adaptation in *T. harzianum*. To our knowledge, such findings have not been previously reported in the context of cellulose degradation for this species. Overall, these results offer valuable insights into how *T. harzianum* responds to environmental changes, revealing a complex regulatory network that is not only crucial for optimizing fungal growth in industrial applications but also deepens our understanding of its biology and ecological interactions.

**Highlights:**

▪ *Trichoderma harzianum* is a key candidate for biotechnology applications.
▪ Light and carbon control CAZymes, photoreceptors, key regulators, and signaling.
▪ *T. harzianum* has distinct gene expression patterns under different light conditions and carbon sources.
▪ *env1*, *cre1*, and *blr2* regulate metabolic adaptation in *T. harzianum*.

**Graphical abstract:** 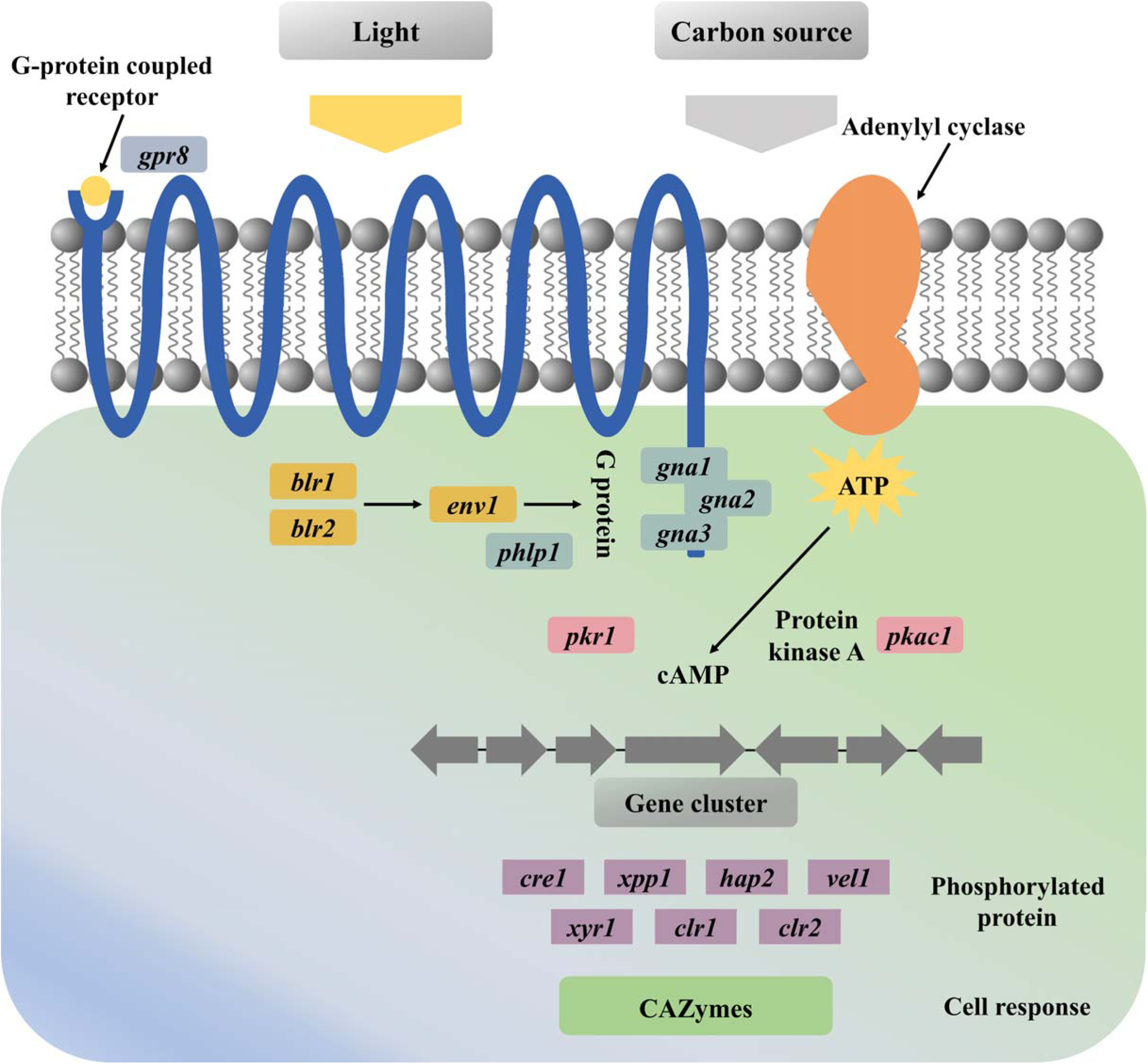

## 1. Introduction

Among *Trichoderma* species, *Trichoderma reesei* is the most extensively studied because of its ability to degrade plant cell walls. It is widely recognized as a model organism for investigating the hydrolytic process during biodegradation (Bischof et al., 2016). Recent research has highlighted the biotechnological potential of *Trichoderma harzianum* (Almeida et al., 2021; Delabona et al., 2020; Delabona et al., 2021; Horta et al., 2018; Liu et al., 2024; Rosolen et al., 2023; Wang et al., 2024) for the exploration of carbohydrate-active enzymes (CAZymes) (Lombard et al., 2014), which are essential genes involved in the breakdown of lignocellulosic substrates (Andlar et al., 2018).

The expression of genes encoding hydrolytic enzymes in *Trichoderma* species is carefully controlled by transcription factors (TFs) (Benocci et al., 2017). The presence of specific substrates stimulates enzyme production, primarily through the Zn_2_Cys_6_-type TF known as xylanase regulator 1 (XYR1) (Stricker et al., 2006). In contrast, easily metabolized carbon sources inhibit this expression via carbon catabolite repression (CCR), which is mediated by the C_2_H_2_-type TF named carbon catabolite repressor 1 (CRE1) (Portnoy et al., 2011). The diverse strategies for cellulose degradation observed in different strains of the same species seem to be linked to the roles of XYR1 and CRE1 (Rosolen et al., 2022). Additionally, other TFs, such as CLR1 (cellulase regulator 1), CLR2 (cellulase regulator 2), ACE1 (activator of cellulase expression 1), ACE2 (activator of cellulase expression 2), ACE3 (activator of cellulase expression 3), HAP2/3/5 (heme activator protein complex 2/3/5), and RXE1 (regulator of xylanase expression 1), act as activators, whereas ACE1 (activator of cellulase expression 1), RCE1 (repressor of cellulase expression 1), and XPP1 (xylanase promoter-binding protein 1) function as repressors in regulating the expression of CAZymes (Aro et al., 2003; Aro et al., 2001; Derntl et al., 2015a; Wang et al., 2019; Zeilinger et al., 2001; Zhang et al., 2019).

In addition to TFs, recent studies have emphasized the role of light as a crucial environmental factor influencing hydrolytic enzyme regulation in *Trichoderma*, with photoreceptors playing a key role (Schmoll, 2018). Orthologs of the photoreceptor genes *wc-1* (*blr1*) and *wc-2* (*blr2*) from *Neurospora crassa* (Benocci et al., 2017), a model organism extensively studied for its photoreception mechanisms and circadian rhythms (Heintzen and Liu, 2007), have been identified in *T. reesei* (Castellanos et al., 2010). In such species, both types of photoreceptors positively regulate cellulase gene transcription in response to light (Stappler et al., 2017). Additionally, the protein ENVOY, similar to the *N. crassa* photoreceptor VIVID (Heintzen and Liu, 2007), modulates cellulase expression in a light-dependent manner (Schmoll et al., 2010). In this context, signal transduction pathways have emerged as crucial for understanding the regulation of hydrolytic enzyme expression (Schmoll, 2018). Notably, in *T. reesei*, both the heterotrimeric G protein pathway and the cyclic adenosine monophosphate (cAMP) signaling pathway are essential for linking light responses with cellulase expression and influencing posttranscriptional regulation (Schmoll, 2018). Connections between regulatory pathways are established mainly via the photoreceptor ENV1 (Tisch et al., 2011).

In the *Trichoderma* genus, research on the influence of light on CAZymes and cellulose-degrading proteins has focused predominantly on *T. reesei*. However, few studies have examined how light affects other metabolic processes within this genus. For example, in *T. atroviride*, light plays a central role in controlling conidiation through the interaction of the photoreceptors BLR1 and BLR2 with mitogen-activated protein kinases (MAPKs), the cAMP signaling pathway, and G proteins (Carreras-Villaseñor et al., 2012; Casas-Flores et al., 2004; Steyaert et al., 2010). These pathways integrate environmental signals and nutritional status, regulating gene expression and conidiation to ensure proper development under various light conditions. Additionally, *blr1* and *blr2* are required for the expression of the light-responsive *phr1* gene, which encodes photolyase (Casas-Flores et al., 2004).

While studies have reported the influence of light on conidiation and the underlying genetic and metabolic processes in *T. atroviride*, there is limited research on *T. harzianum* (Berrocal-Tito et al., 2000), particularly regarding the impact of light on the expression of CAZymes and cellulose-degrading proteins, to our knowledge. Therefore, in this study, we evaluated the regulation of gene expression by CAZymes, TFs, photoreceptors, and proteins related to the heterotrimeric G protein pathway and the cAMP signaling pathway in *T. harzianum*. Specifically, we investigated how different carbon sources and light conditions affect the expression of these target genes. By comparing gene expression under induced or repressed carbon sources and under constant light versus constant darkness, we aimed to analyze the regulatory effects of these environmental factors on gene expression.

The findings revealed distinct patterns of relative expression for genes related to photoreception, TFs, the cAMP signaling pathway, the heterotrimeric G-protein pathway, and CAZymes, which are influenced by light conditions and carbon sources. Notably, our results highlight the significant roles of *env1*, *cre1*, and *blr2* in modulating responses to these environmental factors, with *env1* showing substantial downregulation in darkness, indicating its involvement in light-dependent signaling. In contrast, *cre1* and *blr2* were more strongly influenced by carbon source availability. These findings suggest coordinated regulation between light and nutrient signaling pathways, which is essential for optimizing metabolic responses in *T. harzianum*. Understanding these mechanisms could advance fungal biology, ecology, and industrial applications, where optimizing growth conditions is crucial for biomass degradation and metabolite production.

## 2. Materials and methods

### 2.1. Fungal strains and experimental growth conditions

The Th3844 strain was obtained from the Brazilian Collection of Environment and Industry Microorganisms (CBMAI), located at the Pluridisciplinary Center for Chemical, Biological, and Agricultural Research (CPQBA) at the University of Campinas (UNICAMP), Brazil. Its identity was confirmed by CBMAI through phylogenetic analysis of the internal transcribed spacer (ITS) region, the translational elongation factor 1 (*tef1*), and RNA polymerase II (*rpb2*) marker genes, as described in a previous study (Rosolen et al., 2023). The Th3844 strain was cultivated on potato dextrose agar (PDA) at 28°C for 7–10 days and subcultured up to the third generation. Once fully colonized, the spores were harvested by scraping the surface with sterile water to prepare the spore suspension, which was then used as inoculum for fermentation (**Figure 1**). A Neubauer chamber was used to quantify the number of spores.

**Figure 1.**
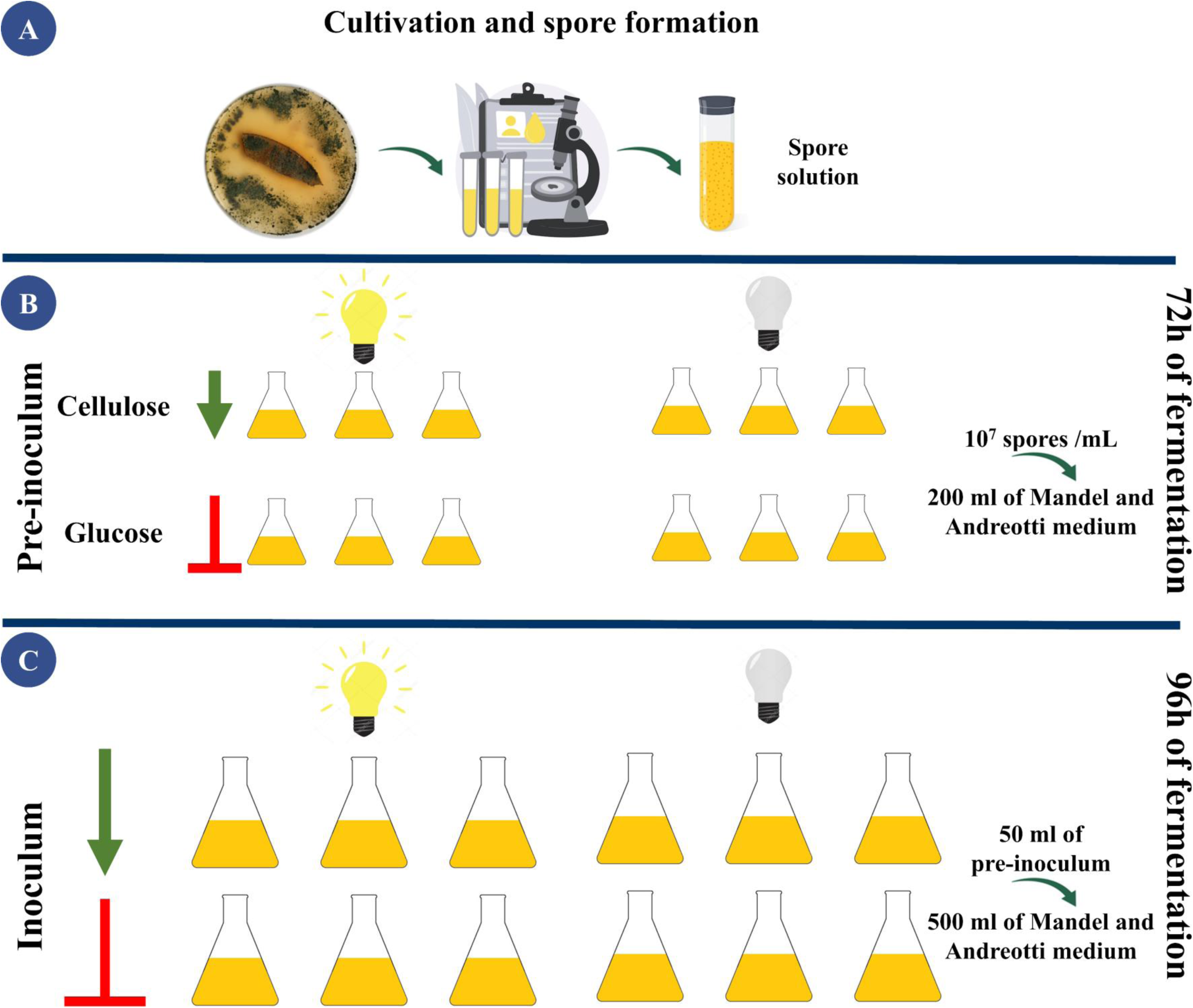
Experimental setup of fermentation with *T. harzianum* using different carbon sources and light conditions. This figure illustrates the experimental setup for fermentation using Th3844 with various carbon sources and light conditions. (**A**) The process began with the cultivation of the Th3844 strain and spore formation, which led to the preparation of the spore mixture used for inoculation. (**B**) In the preinoculum phase, the spores were cultured in media containing either cellulose or glucose as the carbon source under both light and dark conditions. The fermentation volume at this stage was 200 mL, with a spore concentration of 10⁷ spores/mL, and the mixture was incubated for 72 hours. (**C**) The preinoculum was transferred to the final fermentation stage, where 50 mL of the preinoculum was added to 500 mL of media containing cellulose or glucose. Fermentation was continued under light or dark conditions for 96 hours.

Fermentation was performed in biological triplicates, using 200 mL of preinoculum medium, which was inoculated with 10⁷ spores/mL into Mandel and Andreotti media (**Figure 1**). The medium contained either 10 g/L crystalline cellulose or glucose, along with 1 g/L peptone and 100 mL/L mineral base at pH 5.3 (buffered with potassium biphthalate). The mineral base composition was as follows: 20 g/L KH₂PO₄, 14 g/L NH₄SO₄, 3 g/L MgSO₄·7H₂O, 3 g/L CaCl₂·2H₂O, 0.002 g/L CoCl₂, 0.016 g/L MnSO₄·H₂O, 0.014 g/L ZnSO₄·H₂O, 0.05 g/L FeSO₄·7H₂O, and 3 g/L urea. The culture conditions for Th3844 used in this study were established according to Horta et al. (2018).

After 72 hours of incubation under constant light (LED SLIM SMD 30W BIV 6500K IP66 BRONZEARTE) or complete darkness at 28°C on a rotary shaker (200 rpm), 50 mL of the preinoculum was transferred to 450 mL of fermentation medium in a 2 L Erlenmeyer flask (**Figure 1**). The fermentation medium composition varied depending on the carbon source (10 g/L crystalline cellulose or glucose, 1 g/L peptone, 100 mL/L mineral base, 1 mL/L Tween, pH 5.3, buffered with potassium biphthalate). Fermentation proceeded for 96 hours under the same incubation conditions (**Figure 1**). At the end of the cultivation period, mycelia were harvested, rapidly frozen in liquid nitrogen, and stored at −80°C. To prevent interference with gene regulation from light exposure, the harvesting process was conducted under safe red light (Philips PF712E darkroom lamp, red, E27, 15 W). This methodology was based on previous studies that highlighted the influence on gene expression (Beier et al., 2020a; Beier et al., 2020b; Hinterdobler et al., 2020; Hinterdobler et al., 2019; Hitzenhammer et al., 2019).

### 2.2. Isolation of total RNA and cDNA preparation

Total RNA was extracted from 1 mg of each sample using TRIzol Reagent (Invitrogen, Karlsruhe, Germany). The integrity of the RNA was evaluated via 1% (w/v) agarose gel electrophoresis. To assess RNA purity and concentration, a NanoVue spectrophotometer (GE Healthcare, Chicago, IL, USA) was used. For reverse transcription, we utilized the Quantitec Reverse Transcription Kit (Qiagen, Hilden, Germany) to convert the extracted RNA into complementary DNA (cDNA). The synthesized cDNA was diluted at a 1:20 ratio and subsequently stored at −80°C for further analysis using reverse transcription‒quantitative PCR (RT‒qPCR).

### 2.3. Selection of light-responsive genes and conditions for RT‒qPCR analysis

In this study, we selected and designed primers for a total of 20 genes belonging to the following functional categories: photoreceptors, TFs, components of the cAMP pathway, components of the heterotrimeric G-protein pathway, and CAZymes. Specifically, we targeted the following genes: *blr1*, *blr2*, and *env1* for photoreceptors; *clr1*, *clr2*, *cre1*, *hap2*, *vel1*, *xpp1*, and *xyr1* for TFs; *pkac1* and *pkr1* for the cAMP pathway; *gna1*, *gna3*, *gpr8*, and *phlp1* for the heterotrimeric G-protein pathway; and *xyn4*, *egl6*, *bgl1*, and *cel6a* for CAZymes. The coding sequences for each gene were sourced from the genome available in the NCBI database (https://www.ncbi.nlm.nih.gov/datasets/genome/GCA_029844195.1/) and linked to BioProject number PRJNA781962 and the GenBank assembly GCA_029844195.1. The primers were designed using Primer3Plus (Untergasser et al., 2012), aiming for amplicon sizes ranging from 120 to 200 bp, with a target annealing temperature of 60°C and an ideal primer length of 20 bp.

Gene expression was quantified through the continuous monitoring of SYBR Green fluorescence. Each reaction was performed in triplicate, totaling a volume of 8 μL. The components of each reaction included 4 μL of SYBR Green Supermix (Bio-Rad, Hercules, CA, USA), 0.3 μL each of the forward and reverse primers, 2 μL of diluted cDNA, and 1.4 μL of nuclease-free water. Nontemplate controls (NTCs), where cDNA was replaced with sterile water, were incorporated into each primer pair to detect any potential contamination. The reactions were assembled in 384-well plates and carried out using the CFX384 Touch Real-Time PCR Detection System (Bio-Rad, Hercules, CA, USA).

The cycling conditions for qPCR were set as follows: (I) 95°C for 3 minutes, (II) 95°C for 20 seconds, (III) 60°C for 20 seconds, (IV) 72°C for 30 seconds, (V) steps II to IV were repeated for a total of 39 cycles, (VI) a final step at 65°C for 5 seconds, and (VII) a melting curve analysis was performed at 95°C for 5 seconds. The specificity of the amplified products was confirmed through melting curve analysis conducted from 65°C to 95°C. The efficiency of each primer pair was determined by constructing a standard curve using serial dilutions of cDNA (1:10, 1:100, 1:1000, and 1:10000). The amplification efficiency (E) was calculated using the formula *E* = 1 + 10(−1/ *slope*) ^⬚^ *x* 100. The primers were considered effective if their efficiencies fell within the range of 90% to 110%, with R^2^ values between 0.95 and 1.

### 2.4. Measurement of expression levels of selected target genes

The expression variability of the twenty candidate target genes was evaluated by analyzing the variation in their cycle quantification (Cq) values across four experimental groups: (I) light and cellulose (LC, group 1), (II) light and glucose (LG, group 2), (III) dark and cellulose (DC, group 3), and (IV) dark and glucose (DG, group 4) (**Figure 1**). Each experimental group consisted of three biological replicates, resulting in a total of 12 samples (**Figure 1**). Normalization was conducted using the most stable reference gene identified in our preliminary analysis, as detailed in **Supplementary Material 1**. Gene expression levels were calculated using the delta‒ delta cycle threshold method (Livak and Schmittgen, 2001).

To assess whether there were significant differences in gene expression between the control group (LC) and the other experimental groups (LG, DC, and DG), Student’s t test was performed (Student, 1908). However, given that homogeneity of variances between groups was not assumed, Welch’s t test was used (Welch, 1947). To statistically analyze the differentially expressed genes (DEGs), a fold change ≥ 1.5 or ≤ −1.5 and a p value ≤ 0.05 were applied.

## 3. Results

### 3.1. Primer performance analysis

The amplification efficiency (E), correlation coefficient (R²), and slope values of the twenty target genes of Th3844 are presented in **Supplementary Material 1**, which also describes the detailed methodology for the determination of housekeeping genes.

Supplementary Table 2 of Supplementary Material 1 shows the values for the (I) E, (II) R², and (III) slopes as follows: (I) amplification efficiencies from 90.3% (*xpp1*) to 108.7% (*bgl1*); (II) correlation coefficients from 0.952 (*gna3*) to 0.994 (*pkr1*); and (III) slope values from −3.198 (*gpr8*) to −3.130 (*bgl1*). These results demonstrate that the primers designed for all the target genes meet the standards required for RT‒qPCR, making them suitable for subsequent experiments.

Specific fragments of all target genes were successfully amplified from Th3844-derived cDNA using the primers listed in **Supplementary Material 1: Supplementary Table 2**. Each PCR yielded a single product of the expected fragment size, confirming the specificity of the primer amplification. Additionally, the melting curve analysis for each primer pair revealed a single peak, further verifying the primer specificity and ruling out the presence of primer dimers.

### 3.2. Gene expression levels in response to light conditions and carbon sources

The regulation of gene expression is crucial for the adaptation and metabolic response of organisms to changing environmental conditions. In this study, we assessed how different light conditions and carbon sources influence the expression levels of target genes in *T. harzianum*. The RT‒qPCR results revealed significant variations in gene expression across the experimental groups tested, underscoring the complex responses of Th3844 to different light conditions and carbon sources. The Cq values for each gene are shown in **Supplementary Material 1: Supplementary Figures 3-6**.

Gene expression was analyzed through a heatmap with hierarchical clustering, which was generated to illustrate the variation in the expression of different genes under various experimental conditions (**Figure 2**). In general, genes related to the regulation of lignocellulose degradation, such as *xyr1*, *cre1*, *blr1*, and *blr2*, tended to achieve similar expression levels, clustering together on the heatmap (**Figure 2**). Similarly, genes involved in the cAMP signaling pathway and G-protein regulation, such as *gna1*, *gna3*, *pkac1*, and *pkr1*, also clustered, forming another distinct group (**Figure 2**). Moreover, under different conditions, such as LCxLG versus DCxDG, significant changes in gene expression were noted, with certain genes exhibiting markedly increased expression levels (**Figure 2**). Conversely, the expression profiles under the LCxDC and LGxDC conditions were distinct from that under the DCxDG condition (**Figure 2**), indicating that both light and the carbon source play critical roles in regulating gene expression.

**Figure 2.**
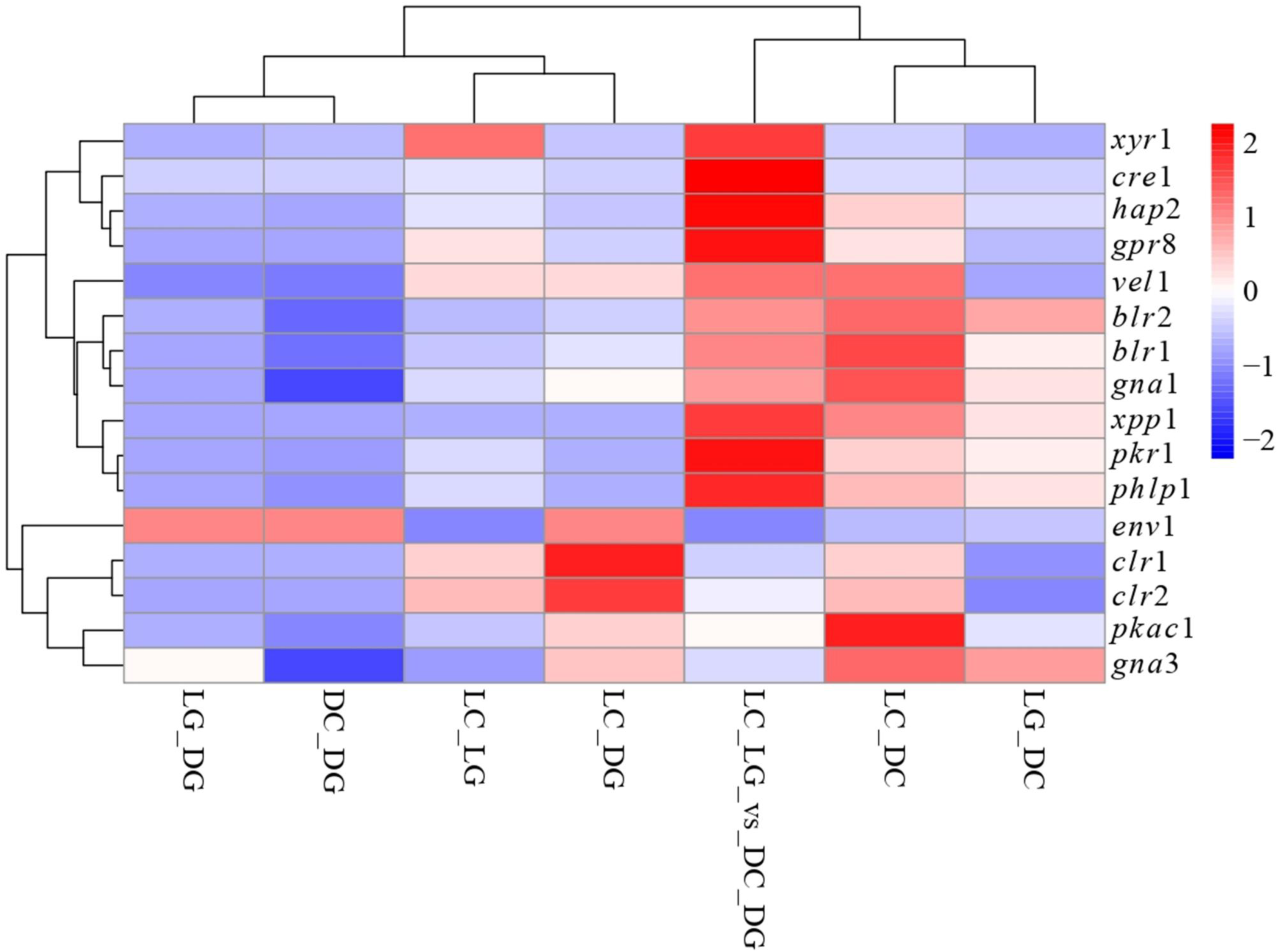
Comparative analysis of gene expression in Th3844 under different light and carbon conditions. The heatmap shows the relative expression levels of key genes across the four experimental groups: light with cellulose (LC), light with glucose (LG), darkness with cellulose (DC), and darkness with glucose (DG). The data represent the ratio of gene expression levels between experimental conditions. The expression values were row-normalized, meaning that for each gene, the values were centered (mean subtraction) and scaled (division by standard deviation) to facilitate comparison across conditions. The colors represent expression levels, with blue indicating low expression, white indicating intermediate expression, and red indicating high expression. Hierarchical clustering was applied to group genes and conditions with similar expression patterns.

More specifically, in class I, which encompasses genes encoding photoreceptors, *blr1* exhibited consistently elevated expression under light and cellulose conditions (**Figure 3**), indicating its crucial role in photoreception and cellulose utilization. Similarly, *blr2* presented increased expression when comparing light and cellulose to both darkness and cellulose and darkness and glucose, whereas *env1* presented lower expression levels when transitioning from light and cellulose to light and glucose (**Figure 3**). These findings suggest that the presence of glucose may suppress *env1* expression when cellulose is available. Conversely, the expression of *env1* was significantly greater in light and cellulose than in darkness, cellulose, darkness, and glucose (**Figure 3**), indicating that light enhances its expression, especially in the presence of cellulose. Notably, its expression decreased significantly under complete darkness, emphasizing its sensitivity to environmental changes.

**Figure 3.**
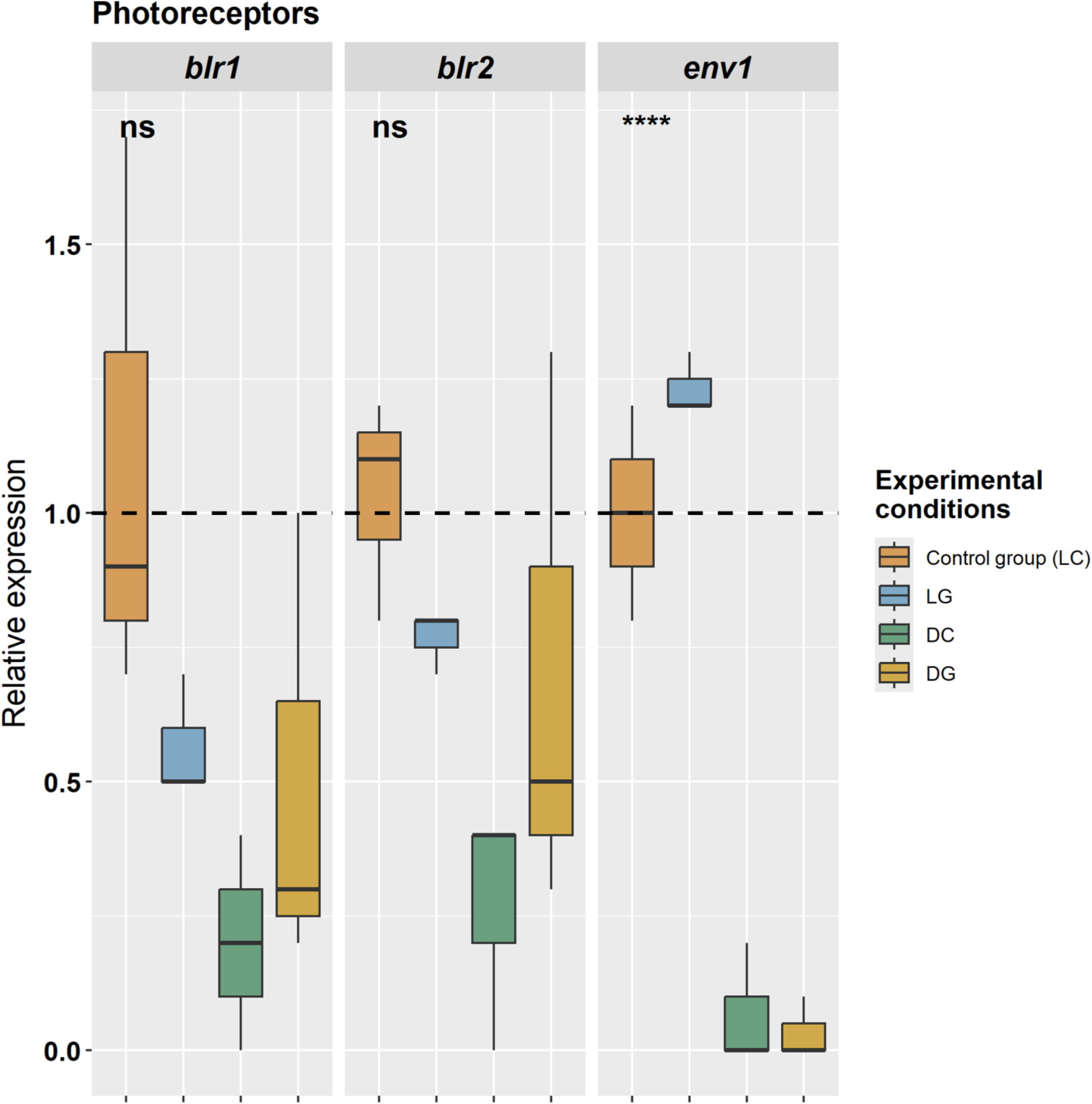
Relative expression of photoreceptors in Th3844 cells determined via qPCR. The relative expression levels of *blr1*, *blr2*, and *env1* in Th3844 were normalized using 18S rRNA and 28S rRNA. The control group (LC) was used as a reference, with a relative expression value of 1. Values above 1 indicate higher expression than the control, whereas values below 1 indicate lower expression. The dotted line indicates a relative expression value of 1. The bars represent the means and standard errors from three biological replicates. Statistical comparisons were performed using ANOVA followed by Tukey’s post hoc test. Asterisks indicate significant differences between groups, whereas "ns" indicates no significant difference. LC: light × cellulose; LG: light × glucose; DC: darkness × cellulose; and DG: darkness × glucose.

In class II, which includes genes encoding TFs, *clr1* consistently presented higher expression in light and cellulose (**Figure 4**), underscoring its pivotal role in regulating responses to cellulose under these conditions. The *clr2* gene also presented increased expression in the presence of light and cellulose compared with the other carbon and light conditions (**Figure 4**), indicating its targeted regulatory function. The expression of *cre1* was notably greater in light and cellulose (**Figure 4**), suggesting its involvement in metabolic pathways favoring cellulose utilization; it exhibited a marked increase in expression in light conditions with cellulose compared with glucose. Additionally, the *hap2* gene displayed significant differences in expression across the experimental groups, whereas *vel1* maintained consistently higher expression under light and cellulose (**Figure 4**), highlighting its central role in transcriptional regulation. *xpp1* presented increased expression in light and cellulose than under other conditions, although its expression was lower in the presence of glucose (**Figure 4**). Finally, *xyr1* was notably expressed in light and cellulose (**Figure 4**), emphasizing its importance in light-specific regulation.

**Figure 4.**
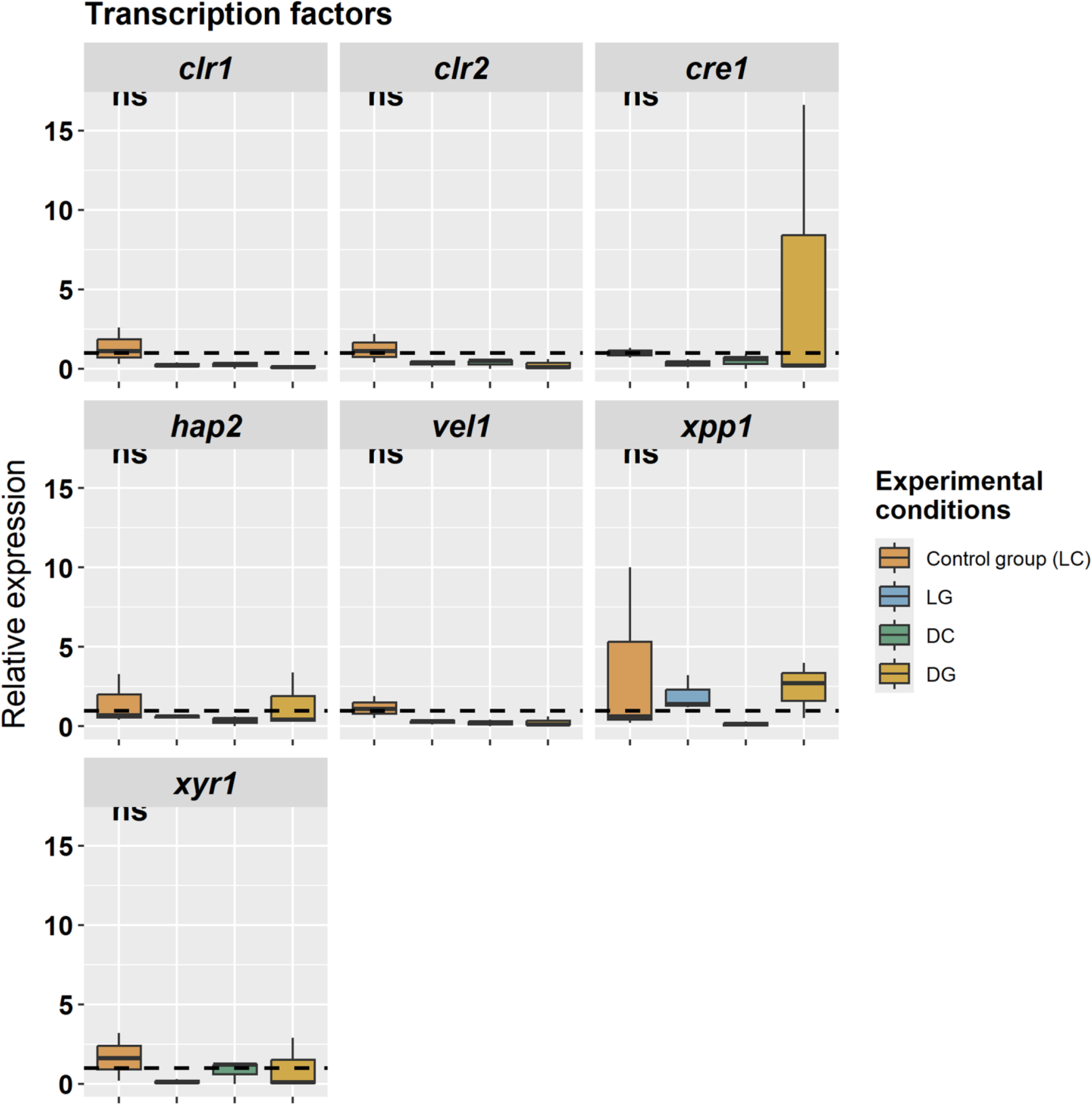
Relative expression of TFs in Th3844 through qPCR. The relative expression levels of *clr1*, *clr2*, *cre1*, *hap2*, *vel1*, *xpp1*, and *xyr1* in Th3844 were normalized using 18S rRNA and 28S rRNA. The control group (LC) was used as a reference, with a relative expression value of 1. Values above 1 indicate higher expression than the control, whereas values below 1 indicate lower expression. The dotted line indicates a relative expression value of 1. The bars represent the means and standard errors from three biological replicates. Statistical comparisons were performed using ANOVA followed by Tukey’s post hoc test. Asterisks indicate significant differences between groups, whereas "ns" indicates no significant difference. LC: light × cellulose; LG: light × glucose; DC: darkness × cellulose; and DG: darkness × glucose.

In class III, which involves genes related to the cAMP signaling pathway, the *pkac1* gene maintained high expression levels across most experimental conditions (**Figure 5**), highlighting its critical role in metabolic regulation under light and cellulose. Overall, *pkac1* presented significantly elevated expression levels under light compared with darkness (**Figure 5**), indicating its potential importance in light-dependent processes. The *pkr1* gene also exhibited high expression but displayed complex regulation (**Figure 5**), suggesting selective responsiveness to specific environmental cues.

**Figure 5.**
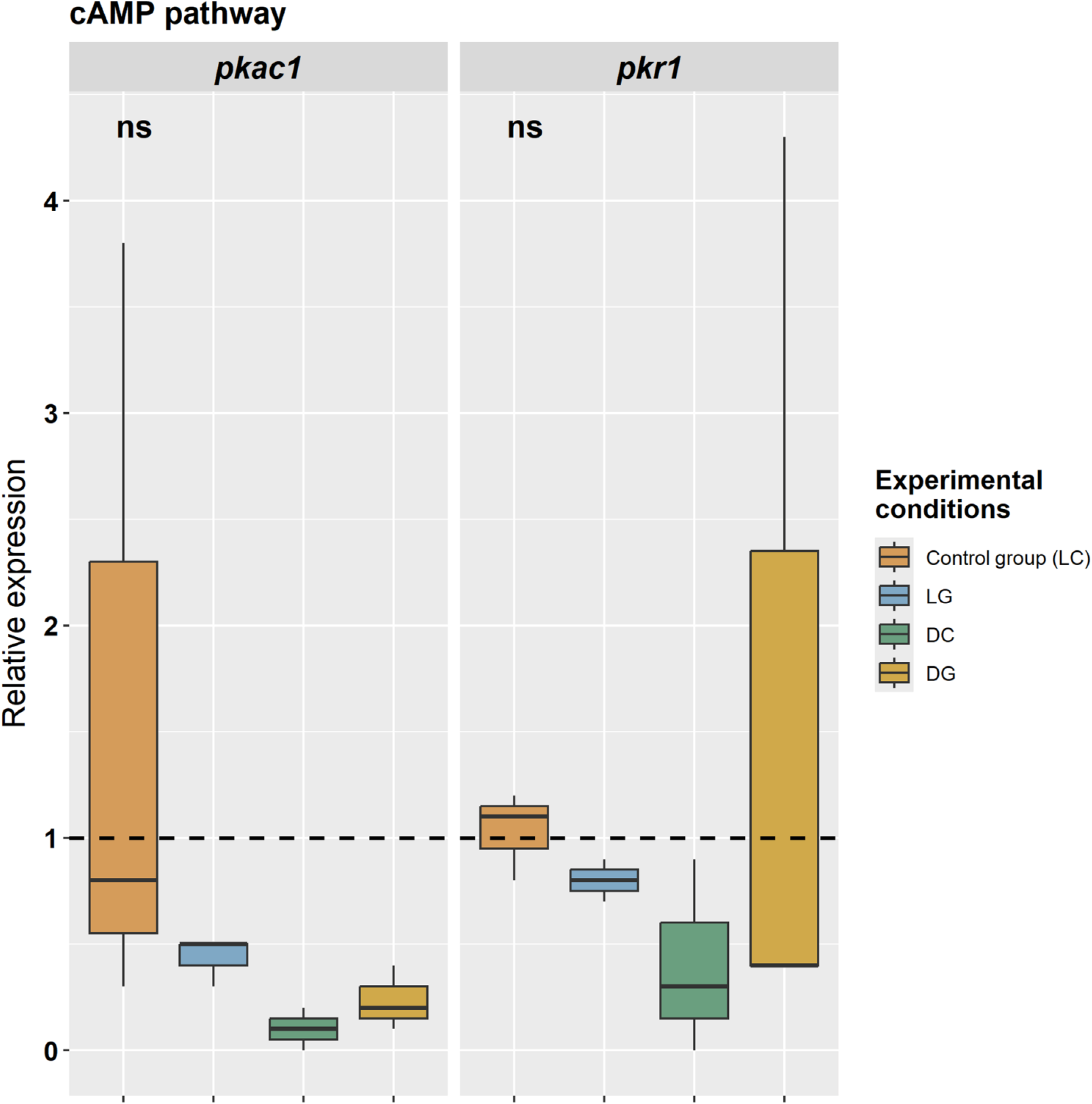
Relative expression of proteins related to cAMP in Th3844 cells determined via qPCR. The relative expression levels of *pkac1* and *pkr1* in Th3844 were normalized using 18S rRNA and 28S rRNA. The control group (LC) was used as a reference, with a relative expression value of 1. Values above 1 indicate higher expression than the control, whereas values below 1 indicate lower expression. The dotted line indicates a relative expression value of 1. The bars represent the means and standard errors from three biological replicates. Statistical comparisons were performed using ANOVA followed by Tukey’s post hoc test. Asterisks indicate significant differences between groups, whereas "ns" indicates no significant difference. LC: light × cellulose; LG: light × glucose; DC: darkness × cellulose; and DG: darkness × glucose.

Class IV, which focused on genes related to the heterotrimeric G-protein pathway, revealed that *gna1* had elevated expression in all comparisons involving light and cellulose (**Figure 3**), indicating its essential role in cellular signaling. The *gna3* gene also presented relatively high expression in several experimental contexts, whereas *gpr8* presented minimal significant differences (**Figure 6**), suggesting that less regulatory influence from environmental factors occurred. Overall, *gna3* presented significantly elevated expression levels under light compared with darkness (**Figure 6**), indicating its potential importance in light-dependent processes. The *phlp1* gene was more highly expressed under light and cellulose conditions than under both darkness and glucose conditions and darkness and cellulose conditions (**Figure 6**), suggesting its potential role in cellulose utilization under light.

**Figure 6.**
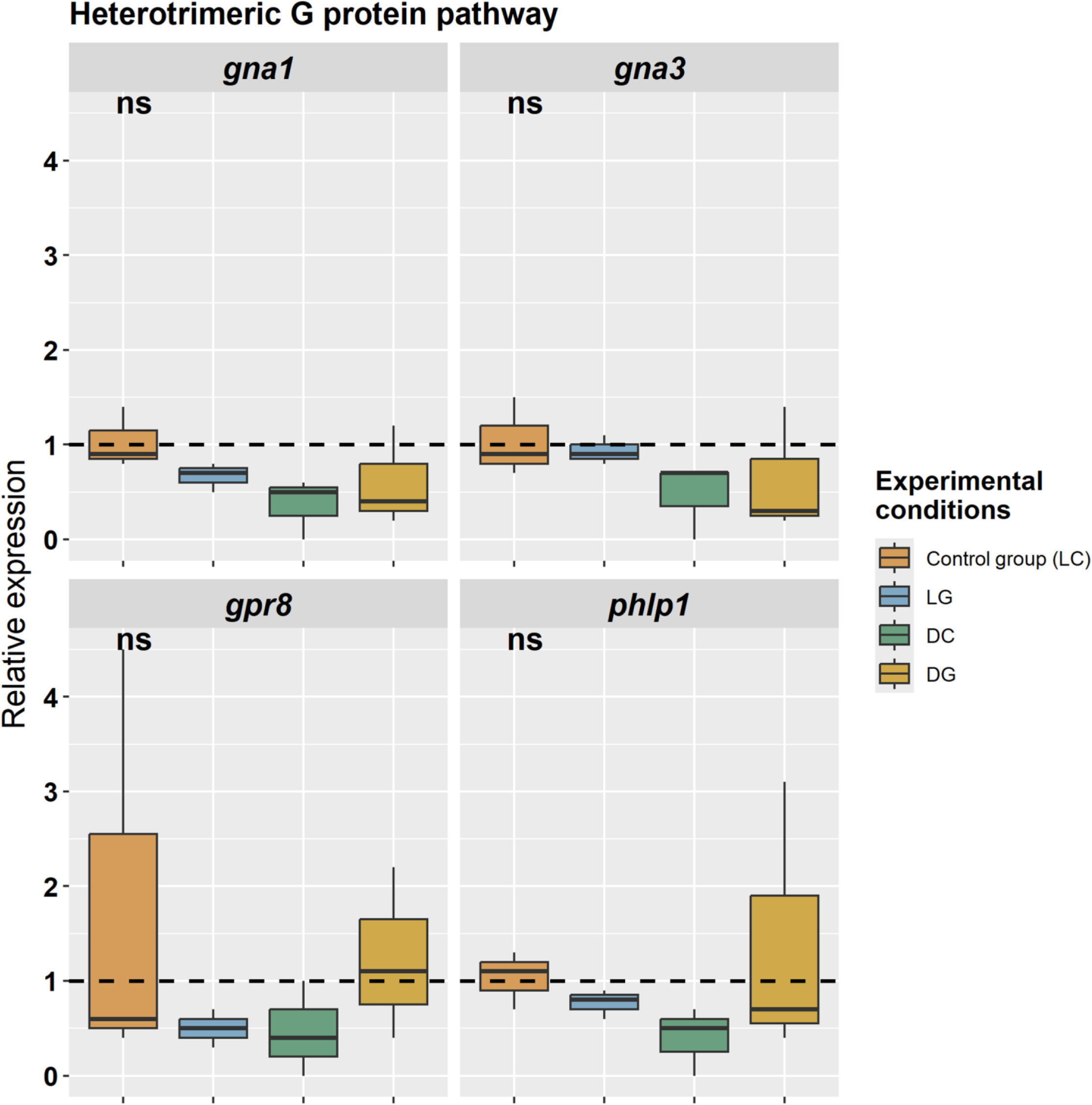
Relative expression of proteins related to heterotrimeric G-protein in Th3844 cells determined via qPCR. The relative expression levels of *gna1*, *gna3*, *gpr8*, and *phlp1* in Th3844 were normalized using 18S rRNA and 28S rRNA. The control group (LC) was used as a reference, with a relative expression value of 1. Values above 1 indicate higher expression than the control, whereas values below 1 indicate lower expression. The dotted line indicates a relative expression value of 1. The bars represent the means and standard errors from three biological replicates. Statistical comparisons were performed using ANOVA followed by Tukey’s post hoc test. Asterisks indicate significant differences between groups, whereas "ns" indicates no significant difference. LC: light × cellulose; LG: light × glucose; DC: darkness × cellulose; and DG: darkness × glucose.

Finally, in class V, which encompasses genes encoding CAZymes, the expression of *xyn4*, *egl6*, *bgl1*, and *cel6a* did not significantly differ across conditions, indicating a stable expression profile regardless of environmental changes.

### 3.3. Analysis of differential gene expression across experimental conditions

Comparative statistical analyses revealed that the expression of most genes did not significantly differ between the control group (light and cellulose) and the other experimental groups (light and glucose, darkness and cellulose, and darkness and glucose). However, three genes displayed statistically significant differences: *blr2* significantly differed between the control group and the darkness and cellulose group (p value = 0.0132) (**Figure 7A**), *cre1* significantly differed between the control group and the light and glucose group (p value = 0.0436) (**Figure 7B**), and *env1* significantly differed between the control group and both the darkness and cellulose group (p value = 0.0048) and darkness and glucose group (p value = 0.0095) (**Figure 7C**). Notably, all three of these genes were downregulated under experimental conditions compared with the control group. For the remaining genes, the p values were not significant (p > 0.05), indicating that the observed differences could be attributed to chance.

**Figure 7.**
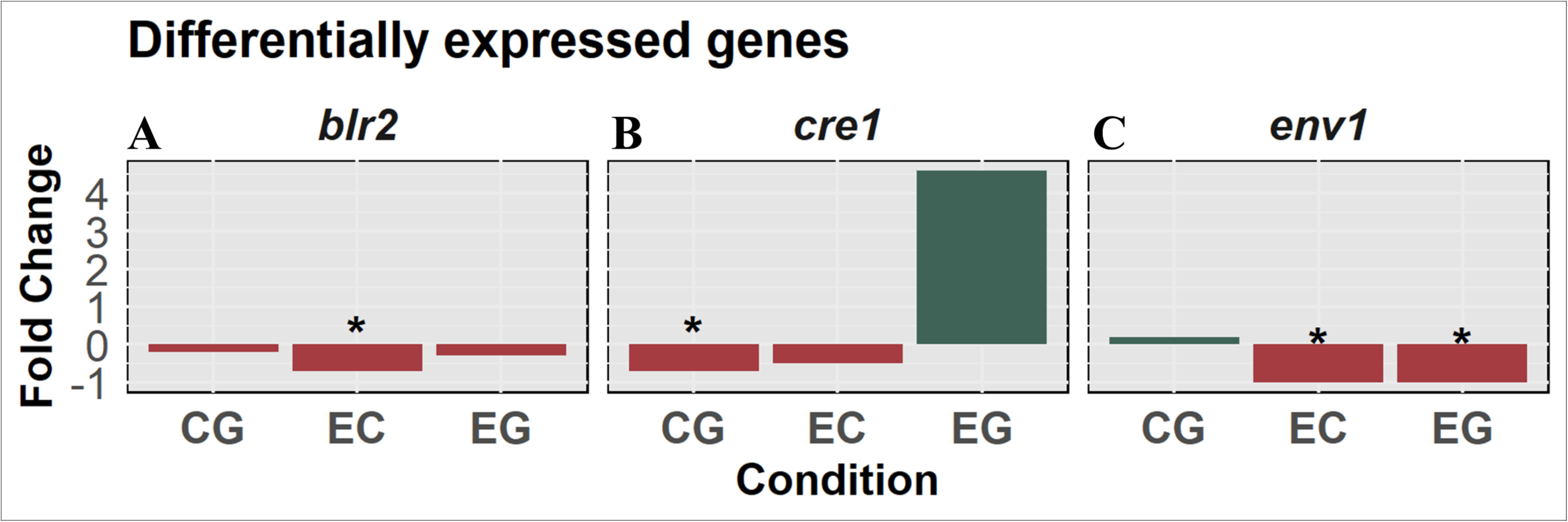
Analysis of differential expression genes (DEGs) in Th3844 under light‒ dark conditions. Comparison of DEGs, including *blr2*, *cre1*, and *env1*, across experimental conditions with LC (light × cellulose) as the control. To statistically analyze the DEGs, a fold change ≥ 1.5 or ≤ −1.5 and a p value ≤ 0.05 were applied. LG: light × glucose; DC: darkness × cellulose; and DG: darkness × glucose. Asterisks reflect the statistical significance of the difference between the control and the tested experimental conditions.

**Figure 8.** Light-mediated regulation of plant cell wall degradation in *Trichoderma harzianum*. The figure illustrates the light-mediated regulation of plant cell wall degradation in *T. harzianum* Th3844 based on mechanistic insights previously reported for *Trichoderma reesei*. The genes investigated in this study are italicized. Not all genes previously described as involved in the light–carbon source response in filamentous fungi are represented. Light perception begins with the G protein-coupled receptor *gpr8*, which detects environmental signals and transmits them to the photoreceptors *env1*, *blr1*, and *blr2*. These photoreceptors regulate downstream signaling pathways, including heterotrimeric G proteins (*gna1*, *gna2*, *gna3*) and *phlp1*, a regulatory component of the G protein signaling pathway. These signaling pathways activate the cAMP cascade, leading to the phosphorylation and modulation of protein kinase A (*pkac1*) and protein kinase R (*pkr1*). These kinases regulate the activity of key transcription factors (*cre1*, *xyr1*, *clr1*, *clr2*, *hap2*, *ve1*, and *xpp1*), which coordinate the expression of genes encoding cellulolytic enzymes. Furthermore, light-regulated genes are organized in clusters within the genome, with several of these clusters overlapping with CAZyme gene clusters. This regulatory network fine-tunes hydrolytic enzyme production, facilitating plant cell wall deconstruction and optimizing carbon acquisition under various environmental conditions.

## 4. Discussion

In fungi, many metabolic processes are regulated by light (Yu et al., 2023; Yu and Fischer, 2019). Specifically, in *T. reesei*, the regulation of cellulase gene expression by light is well documented (Schmoll, 2018). However, to our knowledge, no studies have been reported on *T. harzianum*, particularly on the strain studied in this article. Given the genetic variation among strains within the same genus and the diverse strategies employed by *T. harzianum* strains for cellulose degradation (Almeida et al., 2021; Horta et al., 2018; Rosolen et al., 2022), investigating how abiotic factors influence the expression of genes associated with biomass degradation in this particular strain is essential. Such an investigation is crucial for fully exploring its biotechnological potential. Therefore, herein, we aimed to investigate the relationship between enzyme regulation by carbon sources and light in Th3844 by analyzing the gene expression levels of CAZymes, photoreceptors, TFs, and proteins related to the cAMP pathway and the heterotrimeric G-protein pathway through qPCR. Our two-dimensional analysis included both inducing and repressing carbon sources, allowing us to compare light-specific regulation with dark-specific regulation and eliminating effects associated with a single carbon source.

### 4.1. Gene expression regulation in *T. harzianum* under carbon source and light variations

The observed variations in gene expression indicated that environmental factors, specifically light exposure and the type of carbon source, play a fundamental role in regulating gene expression in *T. harzianum*, particularly in the Th3844 strain. When the effects of the carbon source under both light and darkness were compared, significant differences in gene expression were observed. Under light, LC vs. LG presented notable differences in gene expression levels, whereas in darkness, DC vs. DG presented significant changes in gene expression. These findings suggest a synergistic effect of light and carbon sources on gene regulation, which is not apparent when these factors are analyzed separately.

Furthermore, when the impact of lighting was evaluated while the carbon source was held constant, comparisons between LC and DC and between LG and DG revealed distinct gene expression profiles compared with those observed in the carbon source comparisons under the same lighting conditions. This finding reinforces the idea that both light and carbon sources significantly influence gene activation and repression. Specifically, light seems to act as a crucial regulatory signal in light-sensitive organisms, while the type of available carbon likely modulates the expression of genes involved in carbohydrate metabolism and adaptation to different nutrient sources.

Our results align with those of previous studies showing that environmental factors such as light exposure and carbon sources play crucial roles in regulating gene expression in fungi. For example, research on *T. reesei* has demonstrated that light is a key regulator of developmental and metabolic processes, affecting the expression of genes involved in stress responses and energy metabolism (Beier et al., 2020a; Beier et al., 2020b; Hinterdobler et al., 2019; Hitzenhammer et al., 2019; Schmoll, 2018; Seibel et al., 2009). Similarly, studies on the impact of different carbon sources have shown that nutrient availability modulates gene networks linked to carbohydrate metabolism and secondary metabolite production (Hinterdobler et al., 2020; Hitzenhammer et al., 2019; Missbach et al., 2023; Yu et al., 2023).

Building upon the observed effects of light and carbon sources on gene regulation, we further investigated the role of specific photoreceptors in *T. harzianum* Th3844, particularly the genes *blr1*, *blr2*, and *env1*. These genes have been previously linked to light signaling and metabolic processes (Schmoll, 2018), and our results provide valuable insights into their regulation under different light conditions and carbon sources. Notably, we found that *blr1*, *blr2*, and *env1* presented higher gene expression levels and transcript quantities when the fungus was grown with cellulose as a carbon source than when it was grown with glucose. These findings suggest that these genes may be involved in cellulose metabolism or lignocellulose degradation, which is consistent with findings in the literature indicating that cellulase-related genes are often regulated in the presence of cellulose (Mattam et al., 2022).

In terms of light exposure, our results revealed that *blr1* and *blr2* presented higher gene expression levels under light conditions than under darkness in *T. harzianum* Th3844. These findings support the idea that, in *T. reesei*, BLR proteins may play a significant role in light-responsive processes, potentially regulating light-induced gene expression and cellular responses to light (Castellanos et al., 2010; Schmoll et al., 2010). Additionally, in *T. atroviride*, BLR-1 and BLR-2 are crucial for blue-light responses (Casas-Flores et al., 2004). In the final section of this discussion, the *env1* and *blr2* genes will be further analyzed, with a focus on their roles and potential contributions to industrial biotechnology.

In light of these findings, the expression profiles of TFs in Th3844 under different light and carbon source conditions further illuminate their roles in adaptation to environmental changes. While photoreceptors such as BLR1 and BLR2 mediate light responses, TFs such as CLR1 and CLR2 contribute to the regulation of core cellular functions that are less sensitive to environmental variations. The relatively stable expression levels of *clr1* and *clr2* across all the tested conditions suggest that these TFs may play a fundamental role in maintaining basal cellular functions, regardless of changes in light or carbon source conditions. This stability supports the hypothesis that, in *T. harzianum*, TFs may regulate processes essential for growth and survival, similar to their roles in *T. reesei*, where both CLR-1 and CLR-2 are downregulated by light, yet they do not influence cellulase expression (Beier et al., 2020a; Häkkinen et al., 2014).

In contrast, compared with *clr1* and *clr2*, *cre1* presented higher expression levels across all conditions. This elevated expression highlights *cre1* as a crucial regulatory factor potentially involved in adaptation to environmental changes. The consistent expression pattern of *cre1* across different conditions, with variance primarily observed within biological replicates, suggests a robust role in core regulatory functions, helping the organism respond to diverse environmental stressors in Th3844, as previously reported for *T. reesei* (Campos Antonieto et al., 2016; Han et al., 2020; Portnoy et al., 2011; Rassinger et al., 2018) and *T. harzianum* (Delabona et al., 2021). Like *env1* and *blr2*, *cre1* will be discussed in a separate section. The expression of *hap2* was moderate across all conditions, similar to that of *cre1* but at lower levels. This finding indicates that *hap2* might be involved in regulatory processes that are more condition-specific or finely tuned than *cre1*. In *T. reesei*, HAP2 positively regulates cellulase and hemicellulase gene expression (Benocci et al., 2017). To our knowledge, no studies have reported the influence of light on the gene expression of *hap2* in *T. reesei*.

The expression profile of *vel1* was notably lower during growth on cellulose under light than during growth on glucose and in complete darkness. The similarity in the expression of *vel1* during growth on cellulose and glucose under darkness, as well as on glucose under light, suggests that a conditional regulatory mechanism is influenced by both the carbon source and light. In *Aspergillus nidulans*, VeA coordinates light signals with fungal development and secondary metabolism (Bayram et al., 2008). Its homolog in *T. reesei*, VEL1, is a key regulator of cellulase gene expression (Karimi Aghcheh et al., 2014). However, as these studies were conducted under uncontrolled light conditions, the extent to which this regulation is influenced by light remains unclear. Since VEL1 plays a role in development that varies depending on light conditions and is linked to photoreceptors, cellulase regulation may also be affected by light.

For *xpp1*, relatively high expression levels were observed during growth on cellulose under complete darkness. This elevated expression under specific conditions indicates that *xpp1* may play a significant role in adapting to cellulose utilization in dark environments. In *T. reesei*, XPP1 reportedly regulates the expression of xylanases without affecting cellulase expression (Derntl et al., 2015b; Xu et al., 2023). Additionally, XPP1 plays a role in mediating the transition between primary and secondary fungal metabolism (Derntl et al., 2017). To our knowledge, no studies have reported the influence of light on the gene expression of *xpp1* in *T. reesei*.

Finally, *xyr1* presented a unique pattern, with higher expression during growth on cellulose under light than under other conditions. These findings suggest that, for Th3844, *xyr1* may be involved in metabolic adaptation to cellulose when exposed to light, potentially influencing the expression of genes related to cellulose degradation and light-mediated processes. Overall, these results are consistent with the literature for *T. reesei*, in which the expression level of XYR1 is generally greater when the fungus is grown on cellulose (Furukawa et al., 2009). When *T. reesei* is grown on glucose, the expression of *xyr1* is repressed due to the glucose repression mechanism, which is mediated by regulators such as CRE1 (Castro et al., 2014; dos Santos Castro et al., 2014).

In addition to the regulation of gene expression by photoreceptors and TFs, the cAMP signaling pathway and G-protein coupled receptor signaling also play critical roles in mediating cellular responses to environmental stimuli (Schmoll, 2018). For Th3844, the expression profiles of *pkac1* and *pkr1* under different light conditions and carbon sources reveal insights into their roles in the cAMP signaling pathway and their potential functions in environmental adaptation. The *pkac1* gene, for example, presented higher expression levels under light conditions than under complete darkness. These findings suggest that *pkac1* may play a significant role in the response of the cAMP signaling pathway to light in Th3844. The observed increase in the expression of *pkac1* when grown on cellulose under light conditions and on glucose in complete darkness indicates its involvement in regulating responses to both light and different carbon sources. This differential expression pattern highlights *pkac1* as a key player in adapting the cAMP signaling pathway to environmental changes, potentially influencing metabolic processes and cellular responses on the basis of light exposure and carbon availability. Our findings are consistent with previous research demonstrating that PKAc1 plays a crucial role in the light-dependent regulation of plant cell wall degradation in *T. reesei* (Hinterdobler et al., 2019). This includes involvement in CCR mediated by CRE1, secondary metabolism, and developmental processes (Hinterdobler et al., 2019).

In contrast, *pkr1* presented less pronounced variations in expression levels across light conditions and different carbon sources. The relatively stable expression of *pkr1* suggests that it may play a more constant role within the cAMP signaling pathway, potentially maintaining baseline signaling activity regardless of environmental changes. This stability could imply that *pkr1* is involved in fundamental aspects of the cAMP pathway that are less responsive to variations in light and carbon source conditions. In *T. atroviride*, a previous study suggested that *pkr-1* plays a crucial role in regulating PKA activity and consequently affects the process of asexual reproduction in *T. atroviride* (Casas-Flores et al., 2006). These findings suggest that *pkr-1* influences an organism’s response to light by regulating PKA activity, thereby affecting the process of conidiation and the response to the illuminated environment.

In *T. reesei*, the cAMP signaling pathway is a key component influenced by the heterotrimeric G-protein pathway, with cAMP levels affected by the proteins GNA1 and GNA3 (Schmoll et al., 2005; Schmoll et al., 2009; Seibel et al., 2009; Tisch et al., 2011). Research has highlighted the significant role of the heterotrimeric G-protein signaling pathway in the regulation of cellulase genes, particularly the light-dependent effects associated with the G-alpha subunits GNA1 and GNA3 (Stappler et al., 2017). Compared with those under complete darkness, the expression of *gna1* and *gna3* under light exposure was increased in Th3844, suggesting that light acts as a positive regulatory signal for these genes. This effect could be mediated by light-responsive TFs or signaling pathways that increase gene expression in the presence of light.

For *gna1*, the elevated expression levels under light when grown on cellulose highlight its potential role in cellulose metabolism and suggest that light may increase the expression of genes involved in the degradation of complex carbohydrates. The specific increase in expression under these conditions implies a possible interaction between light and the carbon source in modulating *gna1* activity. This could be due to the activation of light-induced TFs that specifically upregulate *gna1* in response to cellulose.

Conversely, *gna3* exhibited a nearly identical expression pattern between cellulose and glucose, suggesting that its expression is less dependent on the type of carbon source and more uniformly regulated by light. This could imply that *gna3* plays a role in general metabolic processes that are consistently active regardless of the carbon source or that its regulatory mechanisms are not as sensitive to changes in the carbon source as those of *gna1*. In darkness, the reversal of expression profiles for both genes, with increased expression on glucose compared with cellulose, indicates a shift in gene regulation when light is absent. This shift could be reflective of metabolic adaptation where glucose becomes a more prominent energy source in the absence of light, leading to the upregulation of *gna1* and *gna3* in response to glucose availability.

For Th3844, the expression patterns of *gpr8*, a class VII G-protein coupled receptor (GPCR) (Hinterdobler et al., 2020), and *phlp1* under different light and carbon source conditions suggest complex regulatory mechanisms governing gene expression in response to environmental cues. The observation that *gpr8* has similar expression levels under light conditions with cellulose and under dark conditions with glucose indicates that this gene may be involved in pathways responsive to both light availability and carbon sources. In *T. reesei*, the *gpr8* gene plays a critical role in regulating secondary metabolism under dark conditions and is essential for the synthesis of various metabolites (Stappler et al., 2017). Additionally, *gpr8* interacts with the TF YPR2, influencing the activity of the *sor7* gene, which affects both secondary metabolite production and cellulase expression (Stappler et al., 2017).

In contrast, *phlp1* displayed a consistent expression level across different carbon sources under light conditions, indicating that its role in carbon utilization may not be significantly influenced by the type of carbon present. However, *phlp1* was expressed at higher levels on cellulose than on glucose, which suggests that, in Th3844, *phlp1* supports cellulose utilization when light is available. Interestingly, in *T. reesei*, ENV1 and PhLP1 are known to function as reciprocal regulators linking light signaling with nutrient signaling (Tisch et al., 2014). This relationship suggests that *phlp1* could play a similar but less pronounced role in Th3844, which aligns with the notion that *Trichoderma* species may share common regulatory mechanisms.

In our study, we detected undetectable expression of *xyn4*, *egl6*, *bgl1*, and *cel6a* after 96 hours of fermentation with cellulose or glucose as the carbon source under different light conditions. This suggests redundancy among CAZyme genes, with other similar genes potentially compensating for the functions of those that were not expressed. Consequently, these unmeasured genes may not need to be expressed under the given experimental conditions. Another hypothesis is that, after 96 hours of cultivation, CAZymes may have already been expressed and actively functioning on the substrates, even if their gene expression was not detected at that specific time. The dynamics of gene expression and enzymatic activity are fundamental when analyzing the degradation of cellulose and glucose by CAZymes (Corrêa et al., 2020; Huang et al., 2020; Machado et al., 2020). Therefore, the absence of detectable gene expression does not necessarily indicate a lack of enzymatic activity, as the enzymes could still function effectively in the degradation process.

For the cultivation experiments, the 96-hour fermentation period was chosen on the basis of previous studies that indicated high cellulase activity and substantial expression of cellulase and hemicellulase transcripts in the *T. harzianum* strain Th3844 (Almeida et al., 2021; Horta et al., 2018). Thus, this timeframe was deemed appropriate for examining how different carbon sources and light conditions influence the expression of hydrolytic enzymes. Future studies could identify and investigate genes encoding other CAZymes from the genome, and exploring different fermentation time points may provide further insights into enzyme expression and regulation.

Additionally, a previous study using *T. harzianum* as a model indicated that at 96 hours of fermentation, the TFs XYR1 and CRE1 exhibited only basal levels of expression, along with the transcripts coexpressed with both factors (Rosolen et al., 2022). Since the CAZymes investigated are likely regulated by these TFs (Benocci et al., 2017), it can be inferred that the transcript levels of *xyr1* under inducing conditions—specifically, growth on cellulose as the carbon source and light—were insufficient to activate the CAZymes studied in this study. Although *xyr1* expression was higher under the cellulose and light condition than under other conditions, its expression levels were inadequate for effective CAZyme activation in Th3844.

### 4.2. Effects of light and carbon source sources on *env1*, *blr2*, and *cre1*

In this section, we examine the roles of key genes, namely, *env1*, *blr2*, and *cre1,* in how *T. harzianum* responds to specific conditions, namely, the absence and presence of light and the availability of different carbon sources, including cellulose and glucose. Since these genes exhibit differential expression under these conditions, they are discussed individually to emphasize their distinct contributions to metabolic regulation and adaptation strategies.

The *env1* gene is expressed at lower levels in darkness than in light in *T. harzianum*, suggesting its involvement in light-dependent processes, such as the regulation of metabolic pathways influenced by photoreceptors or adaptation to illuminated environments. This reduced expression in the dark, regardless of the carbon source, indicates a possible role in regulating energy metabolism or cellular signaling in response to light. Similar patterns have been observed in *T. reesei* (Castellanos et al., 2010; Schmoll et al., 2005; Schuster et al., 2007). Additionally, in the *T. harzianum* strain Th3844, the pronounced difference in *env1* expression between light and dark conditions suggests a crucial role in low-light adaptation. This finding aligns with previous findings in *T. reesei*, where ENVOY, encoded by *env1*, was involved in light adaptation via a regulatory feedback loop and played a role in nutrient signaling pathways, even in the absence of light (Castellanos et al., 2010; Schmoll et al., 2010; Schuster et al., 2007; Tisch et al., 2011).

Additionally, the downregulation of *blr2* in *T. harzianum* under dark conditions with cellulose (DC) compared with light conditions with cellulose (LC) points to a specific interaction between light and cellulose as a carbon source. This differential expression suggests that *blr2* may play a role in modulating pathways for cellulose utilization in response to light availability. This finding parallels observations in *T. reesei* (Casas-Flores et al., 2004), where BLR2 is involved in light-sensing pathways that regulate carbon metabolism. The reduced *blr2* expression in the dark indicates that *T. harzianum* might employ alternative regulatory mechanisms for cellulose breakdown when light is absent, potentially reflecting an adaptive strategy for efficient resource utilization across different environmental conditions.

In *T. harzianum*, the downregulation of *cre1* in light with glucose, contrary to the expected upregulation observed in *T. reesei* and other filamentous fungi, suggests a distinct regulatory adaptation. Typically, *cre1* upregulation under glucose conditions represses cellulase production as a way to conserve resources when a readily available carbon source is present (de Assis Leandro et al., 2021; Hu et al., 2021; Portnoy et al., 2011). However, the observed downregulation of *cre1* in *T. harzianum* under these conditions might imply that this species has evolved an alternative regulatory mechanism that permits cellulase expression even in the presence of glucose, potentially as a way to remain competitive in environments with fluctuating carbon sources.

This divergence could reflect an ecological adaptation, allowing *T. harzianum* to simultaneously utilize multiple carbon sources or maintain basal cellulase production, possibly facilitating rapid shifts in enzyme expression in response to environmental changes. This suggests that, unlike *T. reesei*, which tightly represses cellulase production in the presence of glucose, *T. harzianum* may integrate glucose and light signals differently, suggesting a unique regulatory network that balances carbon source utilization and environmental responsiveness. Such variations underscore the metabolic versatility of *Trichoderma* species and highlight how *cre1* function may be fine-tuned according to specific ecological contexts.

Overall, the findings regarding *env1*, *blr2*, and *cre1* elucidate a complex regulatory network in *T. harzianum* that enables the organism to adapt to changing environmental conditions. By integrating light and carbon source availability into their metabolic processes, these genes play crucial roles in optimizing both primary and secondary metabolism.

## 5. Conclusion

In conclusion, the comparative analysis between the control and experimental groups of *T. harzianum* highlights the significant influence of light and carbon sources on fungal metabolism and gene regulation. Notably, the differential expression of genes such as *env1*, *blr2*, and *cre1* underscores the intricate interplay between environmental signals and metabolic adaptation. Our findings not only align with previous observations in related fungal systems but also expand our understanding of how *T. harzianum* responds to environmental changes. Moreover, unraveling the regulatory mechanisms behind light-induced gene expression could open new avenues for biotechnological applications, including the optimization of fungal growth for biomass degradation and metabolite production in industrial settings.

## Acknowledgments

We extend our gratitude to CBMAI Campinas and SP for providing the fungal isolates utilized in this study. Our appreciation also goes to the Center of Molecular Biology and Genetic Engineering (CBMEG) at the University of Campinas for granting access to their facilities and laboratory space. We acknowledge the financial support from the São Paulo Research Foundation (FAPESP), the Coordination for the Improvement of Higher Education Personnel (CAPES, Computational Biology Program), and the Brazilian National Council for Technological and Scientific Development (CNPq) for their invaluable contributions to this project.

## Statements and declarations

### Authors’ contributions

**RRR:** Conceptualization; Methodology; Formal analysis and visualization; Writing – original draft. **MACH:** Supervision and Writing – review & editing. **DAS:** Methodology and resources, Writing – review & editing. **APS:** Conceptualization, supervision, review & editing, and funding acquisition.

### Ethics approval

Not applicable.

### Informed consent

Not applicable.

