## Supplementary Material 1 for "Genetic response to light and carbon source variations in *Trichoderma harzianum*: The key regulatory roles of *env1*, *cre1*, and *blr2*"

### 1. Reference genes

#### 1.1. Identifying reference genes for gene expression analysis in *T. harzianum* under varying light conditions

In fungi, genes such as *18S* (18S ribosomal RNA), *28S* (28S ribosomal RNA), *gapdh* (glyceraldehyde-3-phosphate dehydrogenase), *act* (actin), *ef1-α* (elongation factor 1-α), *rpb2* (RNA polymerase subunit 2), *cyp* (cyclophilin), *β-tub* (β-tubulin), and *cyt b* (cytochrome b) are frequently employed as reference genes (Jia et al., 2019; Lian et al., 2014; Singh et al., 2019; Song et al., 2019; Yang et al., 2019; Zhang et al., 2016). However, increasing research indicates that these commonly used housekeeping genes may not maintain stable expression across all experimental conditions (Chapman and Waldenström, 2015). Thus, it is essential to systematically select the most suitable reference genes for specific RT–qPCR experiments in each species to ensure reliable and accurate results.

Although Almeida et al. and Horta et al. (Almeida et al., 2021; Horta et al., 2018) successfully identified and validated reference genes in Th3844 under various carbon source growth conditions, there has yet to be a focused study on the impact of different light conditions on this strain. In this context, we aimed to identify stable reference genes for RT–qPCR-based gene expression analysis in Th3844 cultivated under two distinct light conditions, thereby improving the accuracy and reliability of transcription analysis in this fungus. The assessment of twelve candidate reference genes in fungal cultures subjected to a white light–dark cycle revealed that *18S* rRNA and *28S* rRNA presented the highest stability, making them suitable reference genes for our study.

### 2. Materials and methods

#### 2.1. Reference gene identification and primer development

A selection of twelve reference genes frequently utilized in eukaryotic studies was performed to evaluate stable gene expression in Th3844 cultivated under the specified light conditions. The chosen candidate reference genes included *asl* (ATP synthase lyase γ subunit), *btl* (beta-tubulin gene), *fis1* (encoding a protein involved in mitochondrial membrane fission), *gapdh*, *rho* (GTPase activator), *rpl6e* (ribosomal protein L6e gene), *sar1* (small GTPase of the SAR/ARF type), *tbp* (TATA-binding protein-encoding gene), *tef1* (translation elongation factor 1a gene), and *vma1* (vacuolar ATPase subunit 1 gene), as well as *18S* rRNA and *28S* rRNA. These genes were selected on the basis of reference gene analyses conducted in *T. atroviride* (Flatschacher et al., 2023), *N. crassa* (Cusick et al., 2014), *T. reesei* (Tisch et al., 2011), and *Talaromyces versatilis* (Llanos et al., 2015).

#### 2.2. Evaluation of reference gene expression variability

The expression variability of the twelve tested candidate reference genes was evaluated on the basis of the variation in their cycle quantification (Cq) values across all tested conditions, including (I) light and cellulose (LC), (II) light and glucose (LG), (III) dark and cellulose (DC), and (IV) dark and glucose (DG), in technical triplicate, totaling twelve samples. The analysis was performed considering the mean and standard deviation of the Cq values, with the gene exhibiting the lowest standard deviation identified as the most stably expressed gene, while the gene with the highest standard deviation was recognized as the least stably expressed candidate.

### 3. Results

#### 3.1. Primer performance analysis

Twelve candidate reference genes were selected for Th3844. The amplification efficiency (E), correlation coefficient ($R^2$), and slope values of these genes are presented in **Supplementary Material 1: Table 1.** The values for (I) E, (II) $R^2$, and (III) slope ranged as follows: (I) amplification efficiencies from 90.3% (*btl* and *tbp*) to 106.2% (*vma1*), (II) correlation coefficients from 0.962 (*rpl6e*) to 0.993 (*vma1*), and (III) slope values from -3.578 (*btl*) to -3.182 (*vma1*) (**Supplementary Material 1: Table 1**). These results demonstrate that the primers designed for all 12 candidate reference genes meet the standards required for RT–qPCR, making them suitable for subsequent experiments.

**Supplementary Material 1: Table 1.** Amplification efficiencies, correlation coefficients ($R^2$), and slopes of the RT–qPCR primers for twelve candidate reference genes.

| ***Gene*** | ***Primer forward*** | ***Primer reverse*** | ***Efficiency*** | ***$R^2$*** | ***Slope*** |
|---|---|---|---|---|---|
| ***asl*** | GGCTTCCAACGAGGTCTTCA | GAGACCCTTGTCGGAAGAGC | 94.6% | 0.973 | -3.458 |
| ***btl*** | CACCGTCGTTGAACCCTACA | AGAGCCTCGTTGTCAATGCA | 90.3% | 0.964 | -3.578 |
| ***fis1*** | TACAAGCTCGGCAACTACGG | ACCTTGTCGTCGATGAGCTG | 100.2% | 0.970 | -3.318 |
| ***gapdh*** | GGCATCGTTGAGGGTCTCAT | GGATGATGTTCTGGGCAGCA | 104.2% | 0.992 | -3.226 |
| ***rho*** | CTTGGTTGGCCAGCTGGATA | TGTACGGTCCAAGAACCTGC | 92.2% | 0.977 | -3.524 |
| ***rpl6e*** | TTCCCCTGCGAAGAGTCAAC | GATCTCCTCGATCTTGGCGG | 91.9% | 0.962 | -3.532 |
| ***sar1*** | CAAGCCCGTCGTATCTGGAG | AATCGCTCGTGGTCCTTGG | 104.6% | 0.978 | -3.215 |
| ***tbp*** | TGCCTCTAACGAGCTCACAC | GAGTCACCCCGTTCCCATTT | 90.3% | 0.990 | -3.578 |
| ***tef1*** | GAGAAGATCGACCGCCGTAC | GAAAGCCTCAACGCACATGG | 97% | 0.973 | -3.397 |
| ***vma1*** | CGCCAACAAGAAACAGCGAG | GGACCGGAATCTTGCTTCGA | 106.2% | 0.993 | -3.182 |
| ***18S*** | ATTGGAGGGCAAGTCTGGTG | GGCCCAAGGTTCAACTACGA | 92.4% | 0.990 | -3.518 |
| ***28S*** | TGAAAGGGAAGCGCTTGTGA | GACGAACTGATGCTGGCCTA | 91.9% | 0.977 | -3.532 |

All the candidate reference genes were successfully amplified from the cDNA of the Th3844 strain using the primers specified in Table 1. Each PCR yielded a distinct product matching the anticipated fragment size, confirming the amplification specificity of the primers (data not shown). In addition, the melting curve analysis for each primer pair revealed a single peak, which further substantiates the specificity of the primers and indicates the absence of any primer dimers (**Supplementary Material 1: Supplementary Figure 1**).

**Supplementary Material 1: Supplementary Figure 1.** Melting curves of the twelve candidate reference genes tested (*asl*, *btl*, *fis1*, *gapdh*, *rho*, *rpl6e*, *sar1*, *tbp*, *tef1*, *vma1*, *18S rRNA*, and *28S rRNA*) were generated via RT–qPCR, demonstrating the specificity of the respective primers for *T. harzianum*.

***asl***

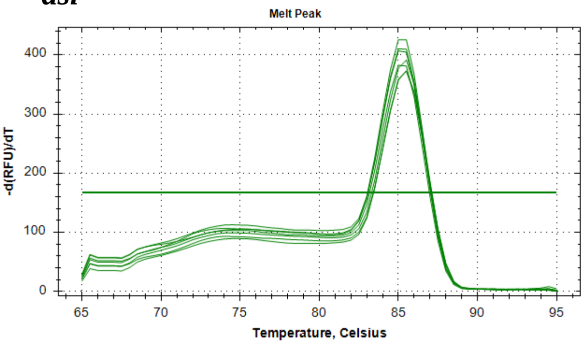


***btl***

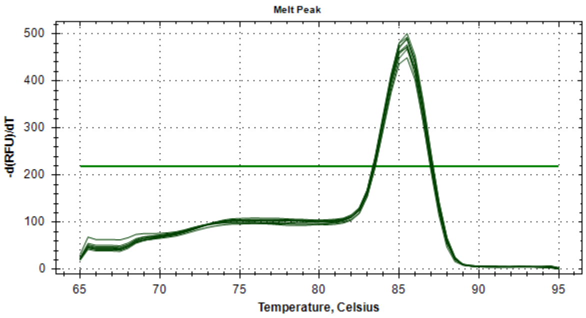

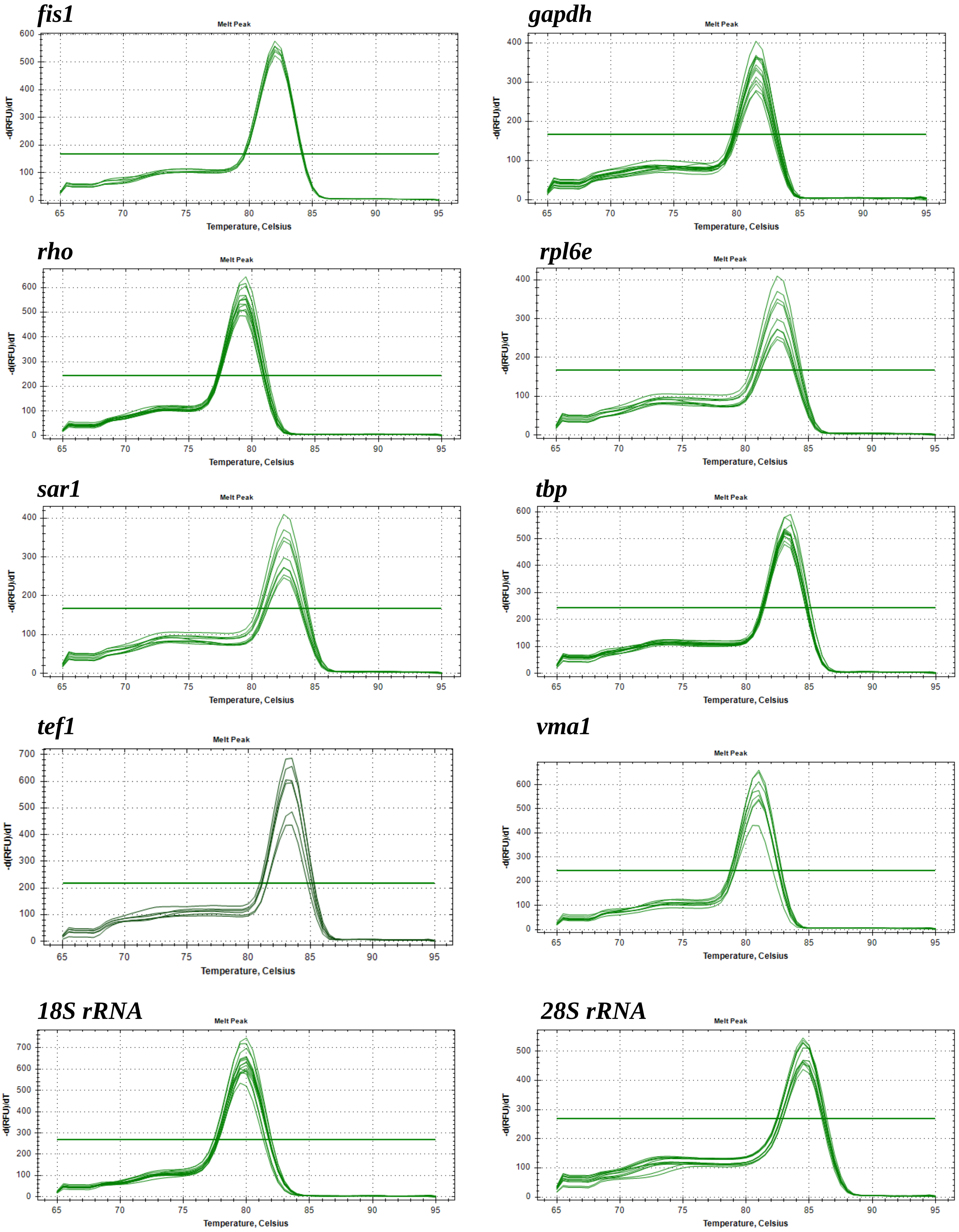


#### 3.2. Variation in the expression of candidate reference genes

The variation in Cq values for the twelve candidate genes was assessed and illustrated using a box plot. This approach facilitated the observation of expression ranges, average Cq values (including both means and medians), and the identification of any outlier values. The average Cq values for the twelve candidate reference genes ranged from 5.6 (*18S*) to 34.0 (*rho*) in samples grown under light or complete darkness, indicating significant variability in expression levels among the assessed genes (**Supplementary Material 1: Supplementary Figure 2**). The average Cq values for the genes *asl*, *btl*, *fis1*, *gapdh*, *rho*, *rpl6e*, *sar1*, *tbp*, *tef1*, and *vma1* displayed notable variation across different experimental conditions. In contrast, the genes *18S* and *28S* presented the least variation in expression levels compared with the other candidate reference genes, making them well suited as reference genes for our analysis.

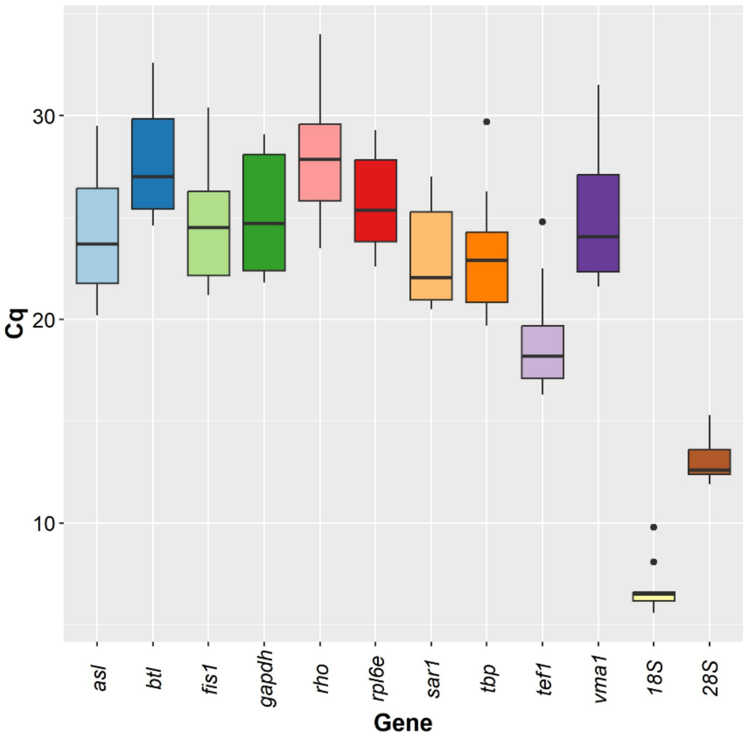


**Supplementary Material 1: Supplementary Figure 2. Evaluation of qPCR Cq values for reference genes in Th3844 under various experimental contexts.** The expression variability of each of the twelve tested reference gene candidates among light–dark conditions in Th3844 is presented as a box-and-whisker plot. Box plots show medians (horizontal lines), 25th to 75th percentiles (boxes), and interquartile ranges (whiskers). The dots indicate the outliers (replicated samples with Cq values above 50% of the interquartile ranges).

### 4. Discussion

#### 4.1. Validation of reference genes for *T. harzianum* under variable conditions

Before analyzing the expression patterns of our genes of interest, we conducted a comprehensive investigation to identify suitable reference genes for our various samples and conditions. On the basis of the literature for *T. reesei*, *T. atroviride*, and *N. crassa* (Beier et al., 2020; Chen et al., 2022; Flatschacher et al., 2023; Hinterdobler et al., 2020; Hitzenhammer et al., 2019; Monroy et al., 2017), we initially selected the genes *asl*, *btl*, *fis1*, *gapdh*, *rho*, *rpl6e*, *sar1*, *tbp*, *tef1*, *vma1*, *18S rRNA*, and *28S rRNA* as potential candidates. However, unlike previous studies, most of these genes presented variable expression levels across our samples and conditions. Although reference genes have been identified for *T. reesei* and *T. atroviride* under different light conditions (Beier et al., 2020; Flatschacher et al., 2023; Hinterdobler et al., 2020; Hitzenhammer et al., 2019; Monroy et al., 2017), these species exhibit significant genetic variation and considerable phylogenetic distance from *T. harzianum* (Rosolen et al., 2022; Rosolen et al., 2023). These factors may have contributed to the differing results observed in our study when these same reference genes were tested for *T. harzianum* under growth conditions similar to those of *T. reesei* and *T. atroviride*. In our study, the most stable reference genes across varying light conditions for Th3844 were *18S rRNA* and *28S rRNA*. These findings are consistent with those of previous studies, which demonstrated that *18S rRNA* and *28S rRNA* are reliable reference genes across various tissues and fungal species (Singh et al., 2019; Xiang et al., 2018; ZHANG et al., 2020).

### Target genes

**Supplementary Material 1: Table 2.** Amplification efficiencies, correlation coefficients ($R^2$), and slopes of the RT–qPCR primers for twenty target genes.

| *Gene* | *Primer forward* | *Primer reverse* | *Efficiency* | $R^2$ | *Slope* |
|---|---|---|---|---|---|
| ***blr1*** | GACCGCAAGCAACGCATTAT | CAGCTCAAACTGCCAGCTTG | 92.2% | 0.954 | -3.524 |
| ***blr2*** | CAATCAGTCGCCCTTTTGCC | TTGTGCTCCAGGAACGAGTC | 102.6% | 0.984 | -3.262 |
| ***env1*** | CAGGAAACGACAGCACTCCT | TTACTCGCAACAGGACACCC | 98.2% | 0.989 | -3.365 |
| ***clr1*** | CTGGTACTCATGCCGCCAA | CATCCAGGTCGTCGACATCC | 107.5% | 0.964 | -3.155 |
| ***clr2*** | TGGAACATGGACATGGACGG | TAATCCACACCCATCGCACC | 95.0% | 0.982 | -3.449 |
| ***cre1*** | GGAAGCTCCCTGTTCCTCAG | CTTCTGCCGCTGTTAGGGAA | 93.1% | 0.982 | -3.501 |
| ***hap2*** | GAGAACTAGATCCCAGCGGC | TGAGGGTCCAGATTCCGACT | 105.4% | 0.977 | -3.198 |
| ***vel1*** | AGTCATCTTTGGCGCCTCTC | GAGGAAGGGGCTCAGTTTGG | 96.2% | 0.988 | -3.417 |
| ***xpp1*** | ATAAAGGTTGAGTCGCCGCA | GGCCGTGTTGTACTGCTTTG | 90.3% | 0.985 | -3.579 |
| ***xyr1*** | TGGGGAGTCTCACTAGCCTC | CGGGGTATCGCAGATCACTC | 101.5% | 0.992 | -3.286 |
| ***pkac1*** | CAGAACCTCCCCAGCATCAG | GAGTTCTGATACGGCGAGGG | 96.7% | 0.965 | -3.403 |
| ***pkr1*** | GGACGAGAGCGATCATCTGG | TAGCTATCGGTGGTAGGGGG | 90.9% | 0.994 | -3.553 |
| ***gna1*** | TGGGTTGCGGAATGTCTACA | TGCATCATCTTGTCCCGCTT | 94.0% | 0.989 | -3.473 |
| ***gna3*** | CGGATGCATGAGCTCCAACA | CCGTTTTGAGTCCTCGTCCA | 95.0% | 0.952 | -3.449 |
| ***gpr8*** | AAGAAGGAATTCGTATCGGTGGA | CGGCGCTCGTATCTCTGG | 105.5% | 0.960 | -3.198 |
| ***phlp1*** | GGAAACCCTCCACCCTGAAG | TCGATTCTGGCGTTGCGATA | 91.0% | 0.984 | -3.557 |
| ***xyn4*** | TGATGTCCACTGTCTCTGTTGT | AGCTTTGTATTGAGAGTTGGTGTTG | 96.8% | 0.984 | -3.402 |
| ***egl6*** | CTCGACTCCAAAAGCCCCAT | CGGGTCAATCTCAAGCGACT | 100.0% | 0.983 | -3.322 |
| ***bgl1*** | CAGCACTGGCAAAACTCACC | GAGACGTGTTTCCAACGCAG | 108.7% | 0.956 | -3.130 |
| ***cel6a*** | CGCTTCTGCAGATGCTTTGC | TACAGGAATGACGGGTTCGC | 100.8% | 0.990 | -3.304 |

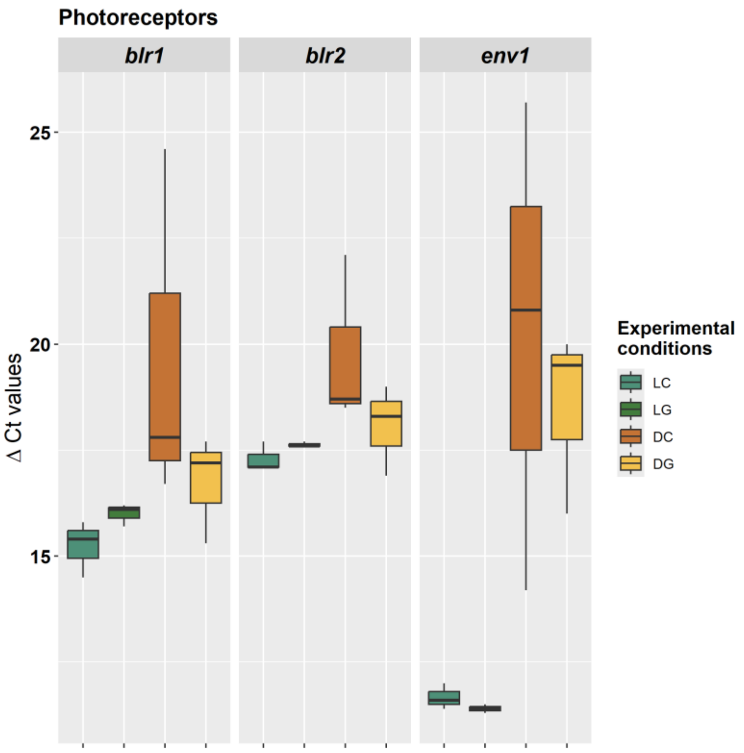


**Supplementary Material 1: Supplementary Figure 3. Variability in the expression of photoreceptors under light–dark conditions in Th3844.** The expression variability of each of the photoreceptors under light–dark conditions in Th3844 is presented as a box-and-whisker plot. The line across the box indicates the median value. The box indicates the 25th and 75th percentiles, and the whiskers represent the maximum and minimum values. LC: light × cellulose; LG: light × glucose; DC: darkness × cellulose; and DG: darkness × glucose.

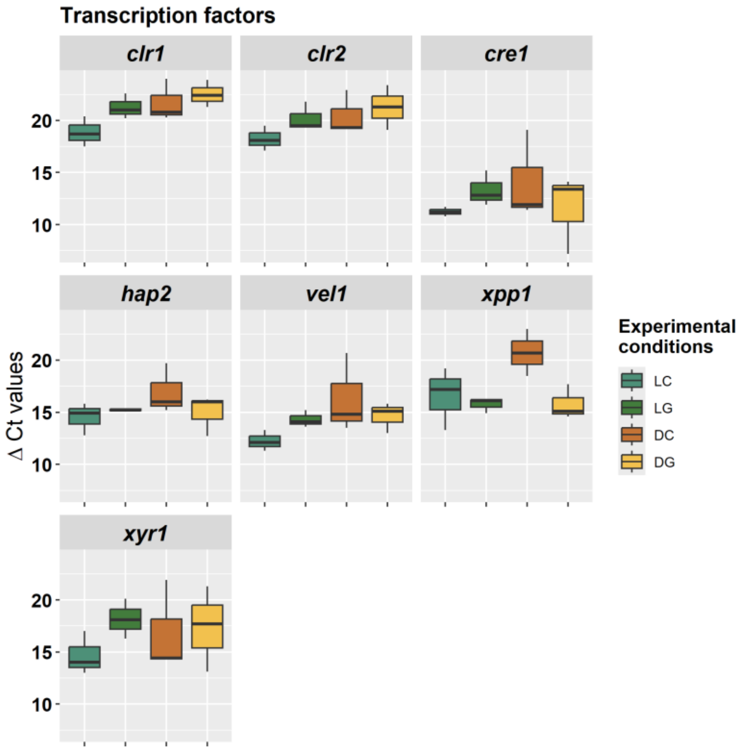


**Supplementary Material 1: Supplementary Figure 4. Variability in the expression of transcription factors (TFs) under light–dark conditions in Th3844.** The expression variability of each of the TFs under light–dark conditions in Th3844 is presented as a box-and-whisker plot. The line across the box indicates the median value. The box indicates the 25th and 75th percentiles, and the whiskers represent the maximum and minimum values. LC: light × cellulose; LG: light × glucose; DC: darkness × cellulose; and DG: darkness × glucose.

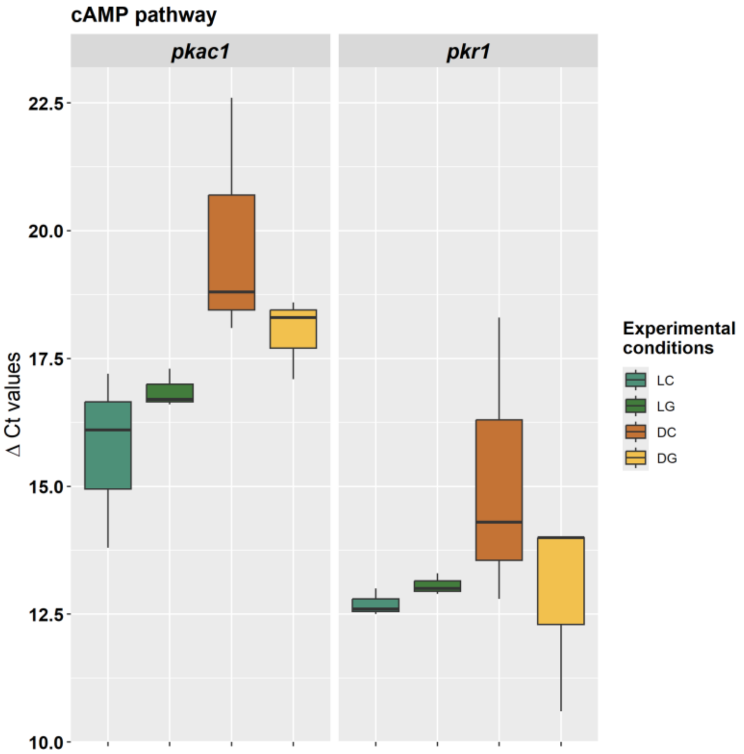


**Supplementary Material 1: Supplementary Figure 5. Variability in the expression of cAMP-related proteins under light–dark conditions in Th3844.** The expression variability of each of the proteins involved in the cAMP pathway under light–dark conditions in Th3844 is presented as a box-and-whisker plot. The line across the box indicates the median value. The box indicates the 25th and 75th percentiles, and the whiskers represent the maximum and minimum values. LC: light × cellulose; LG: light × glucose; DC: darkness × cellulose; and DG: darkness × glucose.

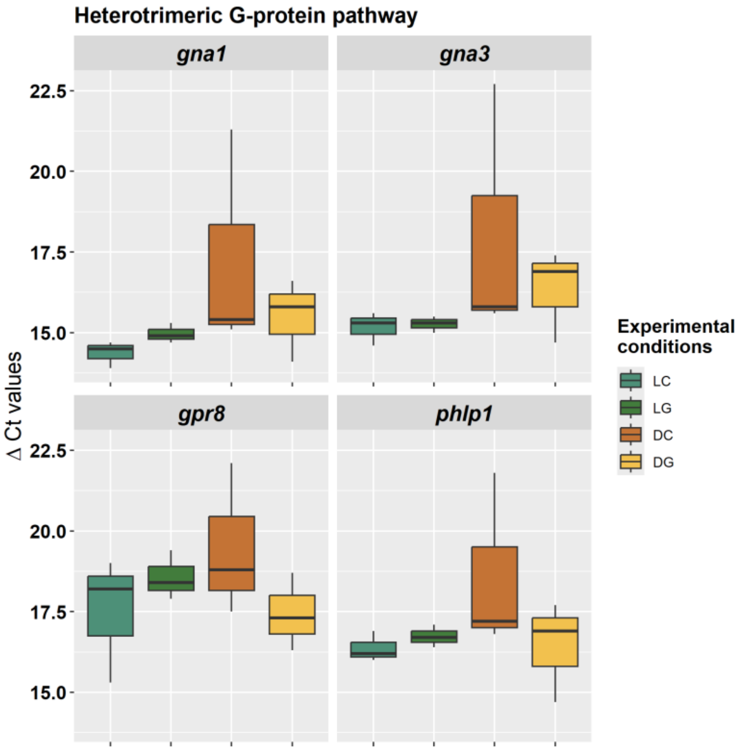


**Supplementary Material 1: Supplementary Figure 6. Variability in the expression of heterotrimeric G protein-related proteins under light–dark conditions in Th3844.** The expression variability of each of the proteins involved in the heterotrimeric G-protein pathway under light–dark conditions in Th3844 is presented as a box-and-whisker plot. The line across the box indicates the median value. The box indicates the 25th and 75th percentiles, and the whiskers represent the maximum and minimum values. LC: light × cellulose; LG: light × glucose; DC: darkness × cellulose; and DG: darkness × glucose.
